# In Silico Therapeutic Intervention on Cytokine Storm in COVID-19

**DOI:** 10.1101/2023.12.05.570280

**Authors:** Abhisek Bakshi, Kaustav Gangopadhyay, Sujit Basak, Amlan Chakrabarti, Abhijit Dasgupta, Rajat K. De

**Affiliations:** Department of Research and Development, Michelin India Technology Centre, Pune, India; Department of Structural Biology, St. Jude Children’s Research Hospital, Memphis, USA; Department of Biochemistry, GITAM (Deemed to be University), Bengaluru, India; A. K. Choudhury School of Information Technology, University of Calcutta, Kolkata, India; Systems Biology Ireland, University College Dublin, Dublin, Ireland; Machine Intelligence Unit, Indian Statistical Institution, Kolkata, India

**Author notes:** Correspondence should be addressed to Rajat K. De, Ph.D., Machine Intelligence Unit, Indian Statistical Institute, 203 B.T. Road, Kolkata, West Bengal 700108, India, or Abhijit Dasgupta, Ph.D., Systems Biology Ireland, University College Dublin, Dublin 4, Ireland,. Equal contribution (Abhisek Bakshi, Kaustav Gangopadhyay, Sujit Basak).

**Keywords:** State space model, PID controller, Cytokines, Kalman filter, ACE2, AT1R, Autoencoder, Molecular docking, Lomefloxacin, and Fostamatinib

## Abstract

The recent global COVID-19 outbreak, attributed by the World Health Organization to the rapid spread of the severe acute respiratory syndrome coronavirus-2 (SARS-CoV-2), underscores the need for an extensive exploration of virological intricacies, fundamental pathophysiology, and immune responses. This investigation is vital to unearth potential therapeutic avenues and preventive strategies. Our study delves into the intricate interaction between SARS-CoV-2 and the immune system, coupled with exploring therapeutic interventions to counteract dysfunctional immune responses like the ‘cytokine storm’ (CS), a driver of disease progression. Understanding these immunological dimensions informs the design of precise multiepitope-targeted peptide vaccines using advanced immunoinformatics and equips us with tools to confront the cytokine storm. Employing a control theory-based approach, we scrutinize the perturbed behavior of key proteins associated with cytokine storm during COVID-19 infection. Our findings support ACE2 activation as a potential drug target for CS control and confirm AT1R inhibition as an alternative strategy. Leveraging deep learning, we identify potential drugs to individually target ACE2 and AT1R, with Lomefloxacin and Fostamatinib emerging as standout options due to their close interaction with ACE2. Their stability within the protein-drug complex suggests superior efficacy among many drugs from our deep-learning analysis. Moreover, there is a significant scope for optimization in fine-tuning protein-drug interactions. Strong binding alone may not be the sole determining factor for potential drugs; precise adjustments are essential. The application of advanced computational power offers novel solutions, circumventing time-consuming lab work. In scenarios necessitating both ACE2 and AT1R targeting, optimal drug combinations can be derived from our analysis of drug-drug interactions, as detailed in the manuscript.

## 1. Introduction

The emergence of novel variants of the severe acute respiratory syndrome coronavirus 2 (SARS-CoV-2) presents a formidable challenge across all sectors of society, notably in healthcare and research and development fields focusing on diagnostics and therapeutic advancements. Initially identified in December 2019, the SARS-CoV-2 rapidly evolved into a global pandemic, manifesting numerous mutations within its genome that have significantly impacted transmissibility, antibody resistance, and disease severity^1^. Recent extensive research has shed light on the interaction between the coronavirus spike protein and human cells, a pivotal step in cellular entry ^2,3^. Yet, due to adaptive mutations in the spike protein gene, the clinical manifestations of SARS-CoV-2 infection remain reminiscent of the earlier patterns, albeit with renewed complexities.

Despite worldwide vaccination efforts, the quest for a lasting solution continues, particularly for immune-compromised individuals and those grappling with persistent post-COVID conditions. Cytokines are central to the intricate web of clinical symptoms and underlying inflammation, pivotal regulators exacerbating the immune response ^4^. A pronounced cytokine surge, dubbed a cytokine storm, has been observed in severe COVID-19 cases, characterized by elevated levels of interleukin-1*β* (IL-1*β*), interleukin-2 (IL-2), interleukin-6 (IL-6), interleukin-17 (IL-17), interleukin-18 (IL-18), and tumor necrosis factor-alpha (TNF-*α*) ^5^. While inflammation initially aids in viral containment, an uncontrolled cytokine storm can inflict harm, necessitating strategies to quell these storms and mitigate patient deterioration.

Our study endeavors to unravel the signaling pathways underpinning cytokine storms, seeking key regulatory nodes as potential drug targets. We employ a control theory-based approach to model these pathways ^6,7^, employing an ordinary differential equation (ODE)-based state-space model (Figure 1) that emulates baseline behavior before the viral assault. Accurate parameter estimation is crucial, and we employ the Kalman filter-based method for approximating these values, acknowledging the intricate relationships within the investigated pathways ^8^. Once the model convincingly mirrors normal pathway dynamics as corroborated by existing literature ^9^, we introduce calibrated perturbations (e.g., elevated ACE2 expressions) inspired by prior research ^10^ to induce a coronavirus-triggered cytokine storm. A proportional-integral-derivative (PID) controller ^6,11^, analogous to a therapeutic agent, scrutinizes the effects of inhibiting/activating specific proteins/molecules (e.g., ACE2 activation or AT1R inhibition in this context) to restore cytokine storm-induced perturbations to baseline pathway dynamics. Our research reveals that ACE2 activation holds great promise as a potential target for drug development. Additionally, our observations indicate that inhibiting AT1R has a comparable impact on the key molecules within this pathway. This comparative study is presented in Figure 5. These findings suggest that AT1R could be a viable alternative drug target for mitigating the cytokine storm.

**Figure 1.**
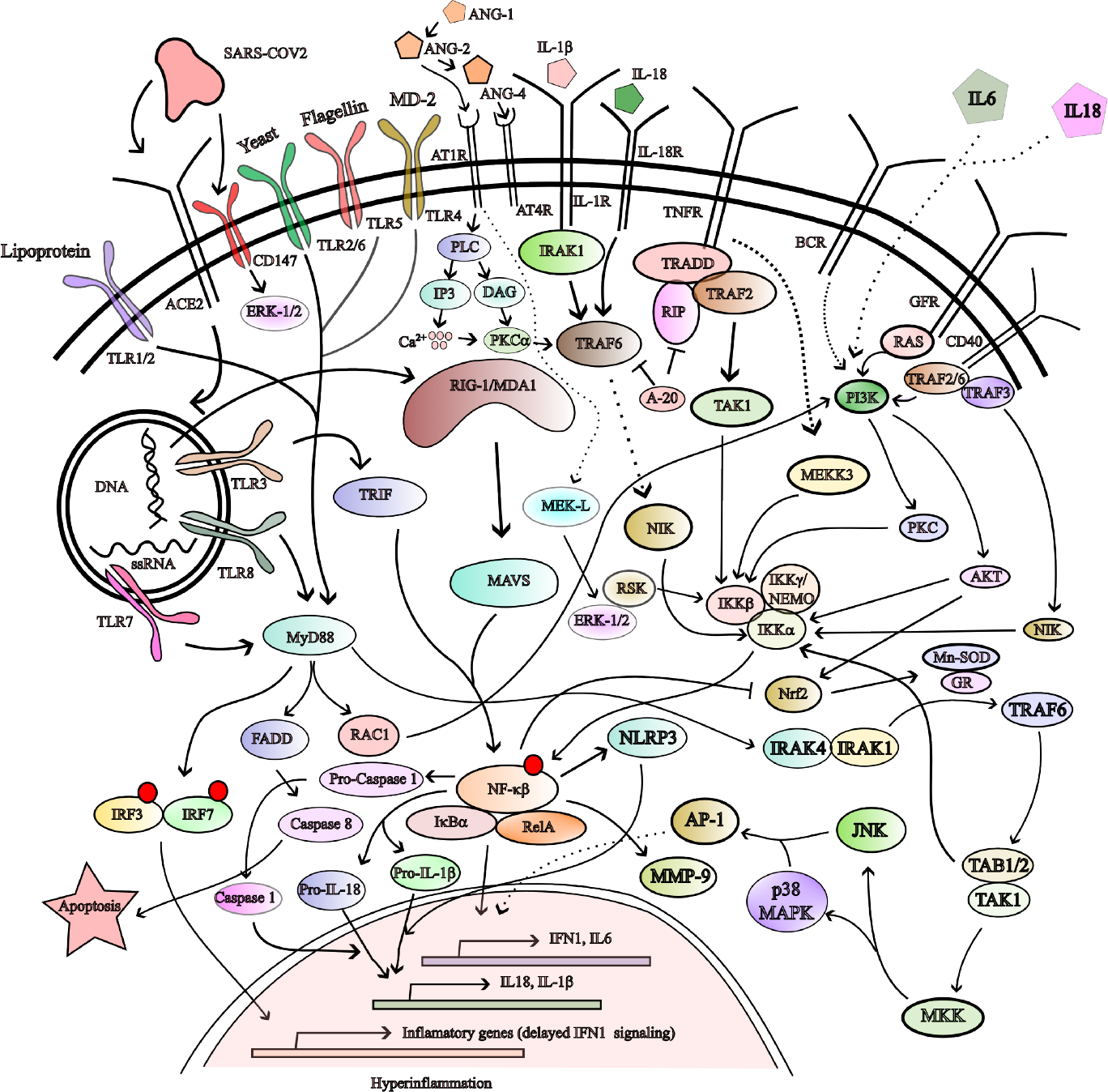
Schematic diagram showing the signaling pathway of cytokine storm by SARS-CoV-2.

Navigating the complexities of a protein-protein network to assess drug efficacy using traditional methods is a time-consuming and financially demanding endeavor, potentially squandering billions of dollars. The scientific community increasingly focuses on deep learning-based drug-target interaction (DTI) predictive models for several compelling reasons. First and foremost, a wealth of comprehensive biological data is readily available in public databases. Additionally, deep learning techniques empower researchers to distill valuable insights from raw data, facilitating the creation of efficient predictive models. In this context, we have harnessed the previous technique ^12^ to identify potential drugs that can individually target ACE2 and AT1R. In scenarios where targeting both ACE2 and AT1R becomes necessary, experts need to anticipate the outcomes of drug-drug interactions. To address this, we have meticulously analyzed the DrugBank database ^13^ to compile a comprehensive list of potential drug interactions with ACE2 and AT1R. These interactions have been categorized as “minor”, “moderate”, and “major”.

We have used a hybrid approach combining deep learning with docking techniques to expedite and streamline drug discovery efforts. Notably, Gentile et al. ^14^ successfully employed deep docking to evaluate numerous drugs against proteins associated with a wide range of diseases. Leveraging this prior knowledge, we have employed a similar methodology to populate our system with well-established drug-protein interaction sites, constructing a sequence-based database rooted in interaction patterns. Furthermore, we have verified drug affinity and its extent using SWISSDOCK, applying these insights to traditional drugs targeted at ACE2 to repurpose them effectively. A parallel investigation was conducted to identify drugs for individual targeting of AT1R. These drugs were categorized based on their affinity into three tiers: low, moderate, and high. Among these drugs, Lomefloxacin and Fostamatinib exhibited exceptional stability when interacting with ACE2 and AT1R proteins, respectively. Furthermore, our analysis of potential drug-drug interactions (DDI) revealed no significant issues when combining Lomefloxacin and Fostamatinib. This observation underscores the safety of administering both drugs concurrently during cytokine storms (CS).

In summary, harnessing advanced computational capabilities to pursue drug repurposing for emerging diseases opens new avenues to explore previously overlooked compounds in the drug bank. This approach equips us to address unforeseen challenges on the horizon with greater confidence and efficiency.

## 2. Materials and methods

This section will talk about the proposed methodology for the task mentioned above. Considering the signaling pathway crucial for cytokine storm (as depicted in Figure 1), we have followed the *in silico* pipeline as depicted in Figure 2.

**Figure 2.**
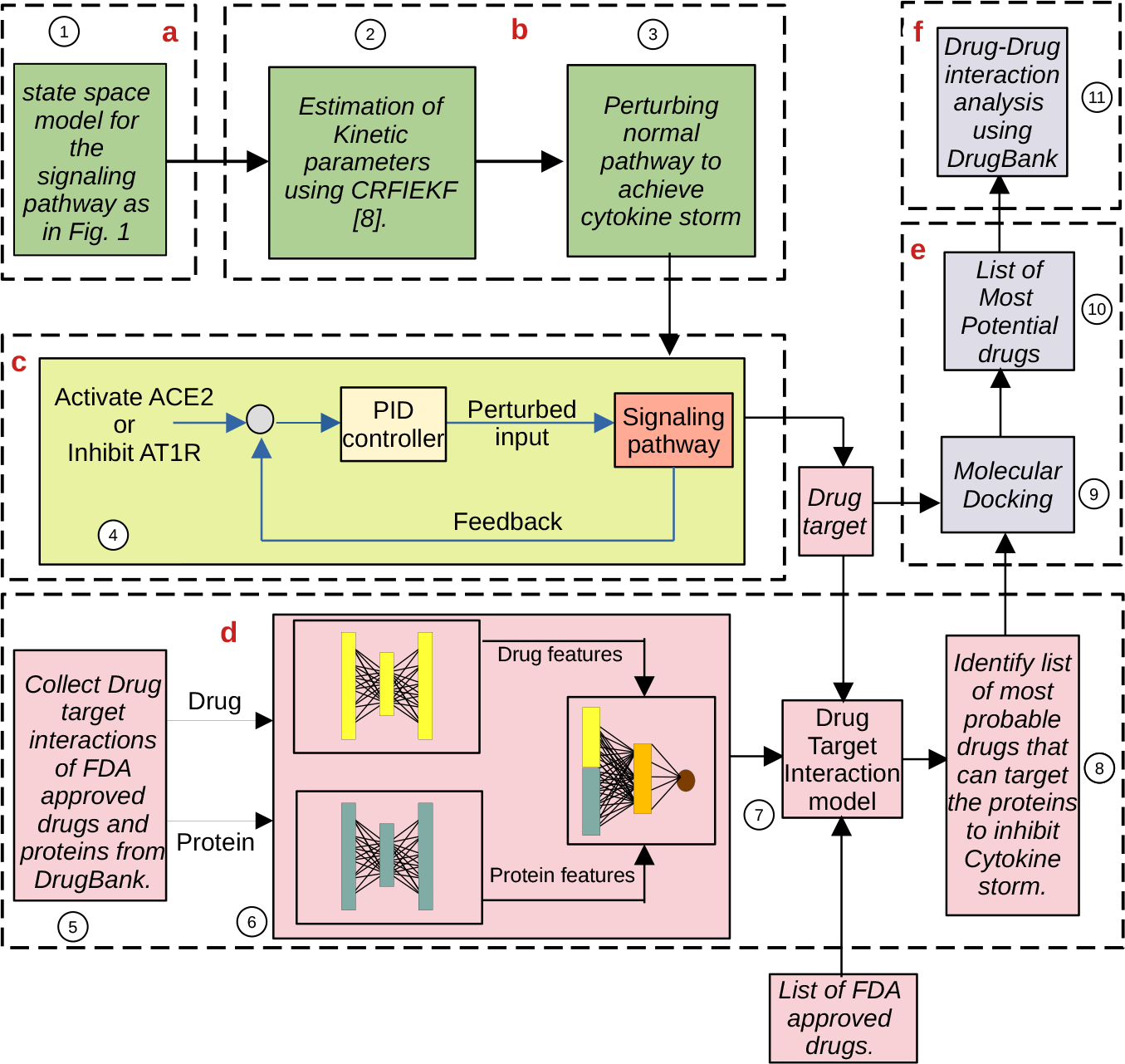
The pipeline of the methodology: a) The formulation of the state space model of the signaling pathway (Figure 1); b) Estimation of kinetic parameters involved in the pathway ^8^ and incorporation of necessary perturbation to meet the state of CS in the pathway; c) Application of PID controller in identifying possible drug target; d) Deep learning model ^12^ for predicting drug-target interaction among FDA approved drugs and target protein(s); e) Molecular docking evaluating the potentiality of the possible drug(s); f) Finding the best probable drugs combination to combat CS.

### 2.1 State space modelling

The following equation represents the state space model of the signaling pathway (Figure 1).

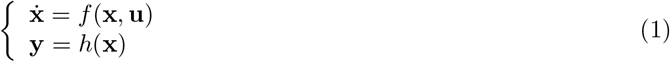

Here, *f* (.) and *h*(.) represent nonlinear functions of state vector **x** (= [*x*_1_, *x*_2_, …, *x*_*n*_]), **y** (= [*y*_1_, *y*_2_, …, *y*_*n*_*′*]) and **u** (= [*u*_1_, *u*_2_, …, *u*_*m*_]) represent expression level of the proteins, end products and external inputs, respectively, involved in the signaling pathway. Let a protein *x*_2_ activates another protein *x*_1_ in the pathway. In addition, an external influence *u*_1_ stimulates/inhibits the activation. Considering the above scenario, the rate of change of *x*_1_ is as follows.

Case 1: Stimulation by *u*_1_

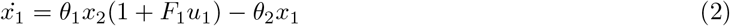

Case 2: Inhibition by *u*_1_

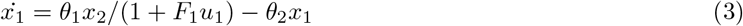

Here, *θ*_1_ and *θ*_2_ are binding rate constant and decay constant, respectively, whereas *F*_1_ represents the feedback constant. This is in line with the investigation carried out by Dasgupta et al. ^11^. The ODEs representing the dynamics of the pathway molecules (Figure 1) are presented in Supplementary Table S1.

### 2.2 Parameter estimation

Numerous kinetic parameters, encompassing binding rates and feedback constants, were derived for the state-space model embodying the specific interest pathway’s inherent behavior. It was accomplished by leveraging the latest technique ^8^. The approach entailed treating these kinetic parameters as the system’s extended state, maintaining their constancy throughout time. A refined adaptation of the Kalman filter methodology was harnessed to generate the measurement signal, drawing solely from the established yet imprecise relationships amongst the molecules within the scrutinized pathway.

These intricate relationships were encapsulated within a fuzzy inference system (FIS), serving as an approximate representation, and a range of estimated parameter values is furnished in Tables 1-2. The imprecise relationship has been exhaustively presented in Supplementary Tables S2-S6. In addition, a comprehensive roster of all estimated parameter values can be found in Supplementary Table S7. The estimation process unfolded iteratively, comprising two distinct phases: “prediction” and “correction”. In the “prediction” phase, an a priori estimation of the state, denoted as 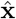, was envisaged, independent of any empirical temporal data. This was sequentially followed by the “correction” step, where 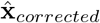 _*corrected*_ was refined based on the established albeit imperfect interrelationships among the pathway molecules. Remarkably, the model adeptly emulated the behavior of the signaling pathway, aligning with earlier inquiries ^15,16^. This fidelity was maintained across parameter values for both unaltered (pre-coronavirus impact) and pathological (cytokine storm) states.

**Table 1.**
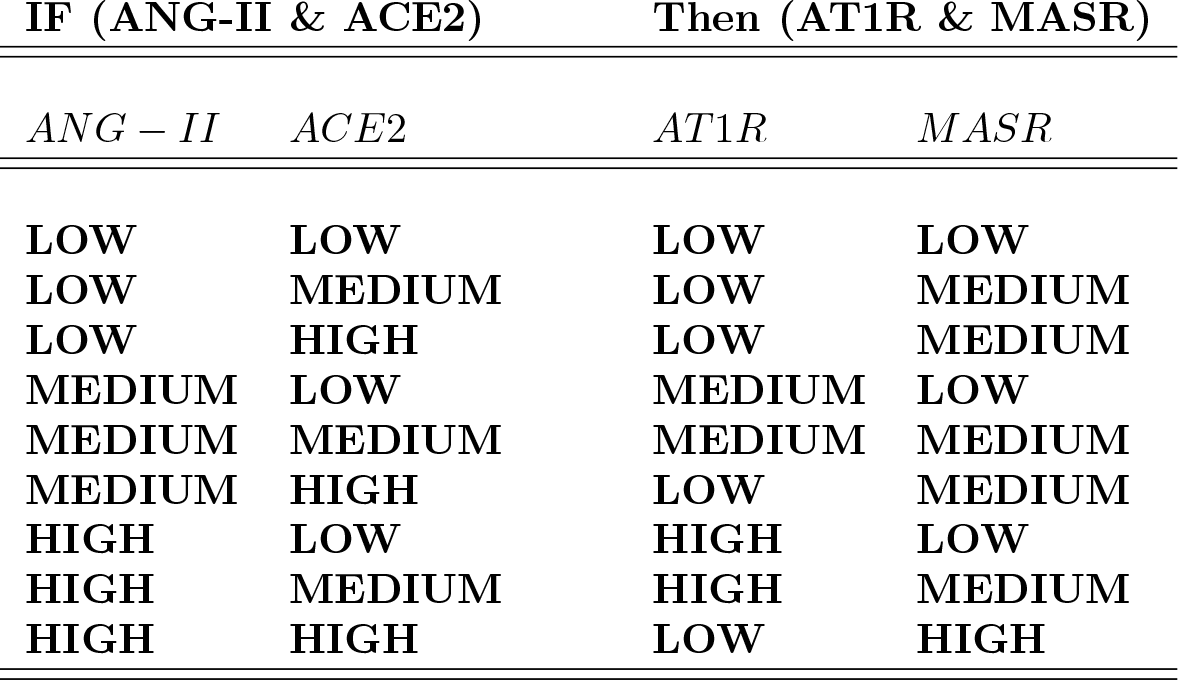
IF-THEN rule base for FIS to capture fuzzy relationship between ANG-II, ACE2 & AT1R, MASR.

**Table 2.**
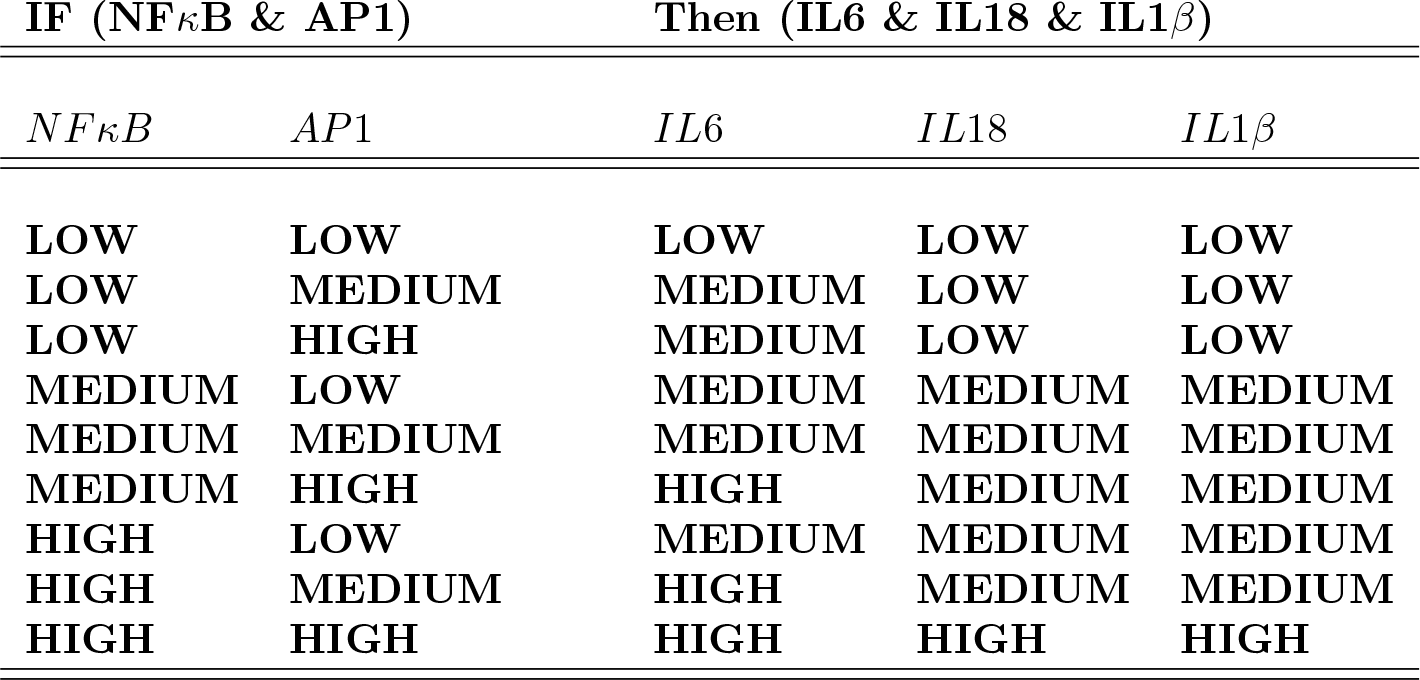
IF-THEN rule base for FIS to capture fuzzy relationship between NF*κ*B, AP1 & IL6, IL18, IL1*β*.

| IF (NFκB & AP1) | | Then (IL6 & IL18 & IL1β) | | |
| --- | --- | --- | --- | --- |
| *NFκB* | *AP1* | *IL6* | *IL18* | *IL1β* |
| LOW | LOW | LOW | LOW | LOW |
| LOW | MEDIUM | MEDIUM | LOW | LOW |
| LOW | HIGH | MEDIUM | LOW | LOW |
| MEDIUM | LOW | MEDIUM | MEDIUM | MEDIUM |
| MEDIUM | MEDIUM | MEDIUM | MEDIUM | MEDIUM |
| MEDIUM | HIGH | HIGH | MEDIUM | MEDIUM |
| HIGH | LOW | MEDIUM | MEDIUM | MEDIUM |
| HIGH | MEDIUM | HIGH | MEDIUM | MEDIUM |
| HIGH | HIGH | HIGH | HIGH | HIGH |

### 2.3 Perturbing the signaling pathway to achieve cytokine storm

A recent study ^17^ unveiled that the ACE2 receptor protein facilitates the cellular entry of SARS-CoV-2. Interaction between the virus and ACE2 prompts the downregulation of ACE2 itself ^18^. Consequently, this interaction triggers an upsurge in angiotensin II via ACE, given the diminished ACE2 presence, which typically converts angiotensin to the vasodilatory angiotensin 1–7^19^. As ACE2 levels decrease, angiotensin I and angiotensin II degradation diminishes, leading to a gradual elevation in their plasma concentrations. An intensive care unit team in the United States underscored the correlation between elevated angiotensin 1–10 and reduced angiotensin 1–9 (a product of ACE2 processing) with unfavorable prognoses in cases of acute respiratory distress syndrome (ARDS) ^20^.

Conversely, following SARS-CoV-2 entry, the downregulation of ACE2 curtails its capacity to break down Ang I and Ang II into Ang-(1-9) and Ang-(1–7), respectively. This situation triggers an excessive activation of the classical ACE-Ang II AT1R axis within the renin-angiotensin system (RAS), marked by vasoconstriction, inflammation, apoptosis, fibrosis, and oxidative stress. Given this intricate interplay, we have adopted low expression values for ACE2 (approximating 0, denoting viral entry) to simulate the conducive conditions for cytokine storm (CS).

### 2.4 Application of a PID controller

Findings from earlier studies suggest that activating ACE2 could serve as a potential avenue for reinstating the normal functioning of the pathway, effectively curtailing the cytokine storm. This compelling notion propelled us to delve deeper into the feasibility of inhibiting AT1R as an alternate strategy to achieve the same outcome. Consequently, we employed two distinct proportional-integral-derivative (PID) controllers, emulating drug-like interventions, to ascertain the effectiveness of either ACE2 activation or AT1R inhibition in resolving the predicament. The simulation was carried out within the Simulink platform of the Matlab software, enabling the nuanced exploration of these interventions.

The reference signal was meticulously set to approach unity, while for ACE2 and AT1R targets, it was pegged close to one and zero, respectively. Employing this framework, the error signals underwent iterative refinement, optimizing the expressions of the various molecules integral to the pathways corresponding to each target. Mathematically, the error signal e(t) at any given time point t can be represented as follows.

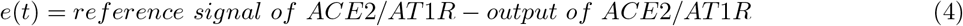

Finally, at time *t, e*(*t*) led to the control signal *c*(*t*) for ACE2 activation/ AT1R inhibition using the following equation,

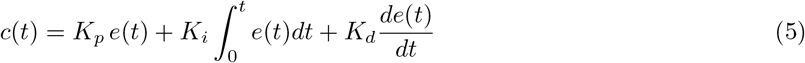

Here, the constant terms *K*_*p*_, *K*_*i*_, and *K*_*d*_ can be termed as proportional, integral, and derivative gains, tuned to particular values to produce the appropriate control signal(s).

In particular, our analysis accounts for SARS-CoV-2 as an external input that influences ACE2 activities. We characterize this influence during normal and infected conditions by attributing a high value (= 0.99) to SARS-CoV-2 for the former and a low value (= 0.01) for the latter. Drawing on the previously mentioned studies, we postulate that directing therapeutic efforts toward ACE2 could hold promise in mitigating this scenario. To evaluate this hypothesis, we deployed a PID controller (with coefficients *K*_*p*_= 0.0028; *K*_*i*_= 5.7991; *K*_*d*_= -0.0036) to regulate the impact of SARAS-CoV-2 on ACE2. The central objective was to validate whether modulating this targeted factor could effectively curtail the excessive production of cytokines.

Conversely, AT1R is implicated in a spectrum of pro-inflammatory activities encompassing vasoconstriction, hypertrophy, fibrosis, and more ^21^. Given that ACE2 activation influences AT1R, we contemplate whether targeting AT1R might offer an alternative to ACE2 intervention. To probe this proposition, we established an independent PID controller (with coefficients *K*_*p*_ = 0.0340, *K*_*i*_ = 0.0159, and *K*_*d*_ = -0.0006) to regulate the impact of SARAS-CoV-2 on AT1R. The overarching aim was to assess whether this configuration could achieve the earlier objective.

### 2.5 Deep learning model for drug-target identification

In this context, a sophisticated deep learning model ^12^ was harnessed to anticipate the potential affinity of drugs with prospective targets, namely ACE2 or AT1R. Leveraging the comprehensive DrugBank dataset, our initial step involved training the model using a diverse assortment of positive and negative drug-target-interaction (DTI) pairs. The DrugBank resource furnishes a collection of FDA-approved drugs and their target proteins. Drawing from this repository, we gathered a dataset encompassing 2505 FDA-approved drugs and 1198 target proteins to establish the foundation of our DTI pairs. It is pertinent to note that DrugBank exclusively cataloged positive DTI relationships. Consequently, to create a robust set of negative samples, we judiciously selected proteins conspicuously absent from the list as potential drug targets. This considered approach transformed the DTI predictive task into a binary classification challenge.

For the numerical representation of both drugs and target proteins, we harnessed the extended connectivity fingerprint (ECFP) technique using the Morgan algorithm ^22^, coupled with protein sequence composition descriptors (PSC) ^23^, respectively. The feature extraction process involved the deployment of two distinct autoencoders, individually tailored to capture the intricate attributes of drugs and proteins. The outcome of this feature extraction phase was subsequently amalgamated and channeled into fully connected classification layers. The sigmoid activation function was embedded in the model’s output layer to facilitate predictive outcomes. Remarkably, this model achieved an impressive accuracy of 89% on an independent test dataset, validated through a rigorous five-fold cross-validation procedure.

### 2.6 Molecular docking

The molecular docking was performed using SWISSDOCK following Grosdidier et al. ^24^. ZINC15 database provided the structure of the small molecules ^25^. Briefly, the SARS-COV2 receptor-binding domain’s (RBD) PDB structure, spike protein (PDB ID: 7DQA), and small molecules were selected from the ZINC15 database. We chose the structures with better ΔG values.

### 2.7 Analysing drug-drug interaction

Having identified distinct candidates capable of targeting ACE2 and AT1R through our deep learning model, as elaborated in the preceding section, our present focus shifts to exploring potential drug-drug interactions (DDIs) within these two separate sets of drugs. This exploration becomes particularly relevant when medical exigencies necessitate the application of distinct drugs to target ACE2 and AT1R individually. We harnessed the approach devised by Dmitriev et al. ^26^ to carry out this investigation. This method classifies the risk levels of DDI into three categories: major, moderate, and minor, providing a comprehensive assessment of the potential interactions.

Additionally, for a more comprehensive understanding of these DDIs, we leveraged the capabilities of the DrugBank ^13^ toolbox. In a concise overview, our analysis unearthed a solitary major interaction, accompanied by seventeen moderate interactions and five minor interactions. The comprehensive summary of these interactions has been meticulously compiled and documented in Table 3.

**Table 3.**
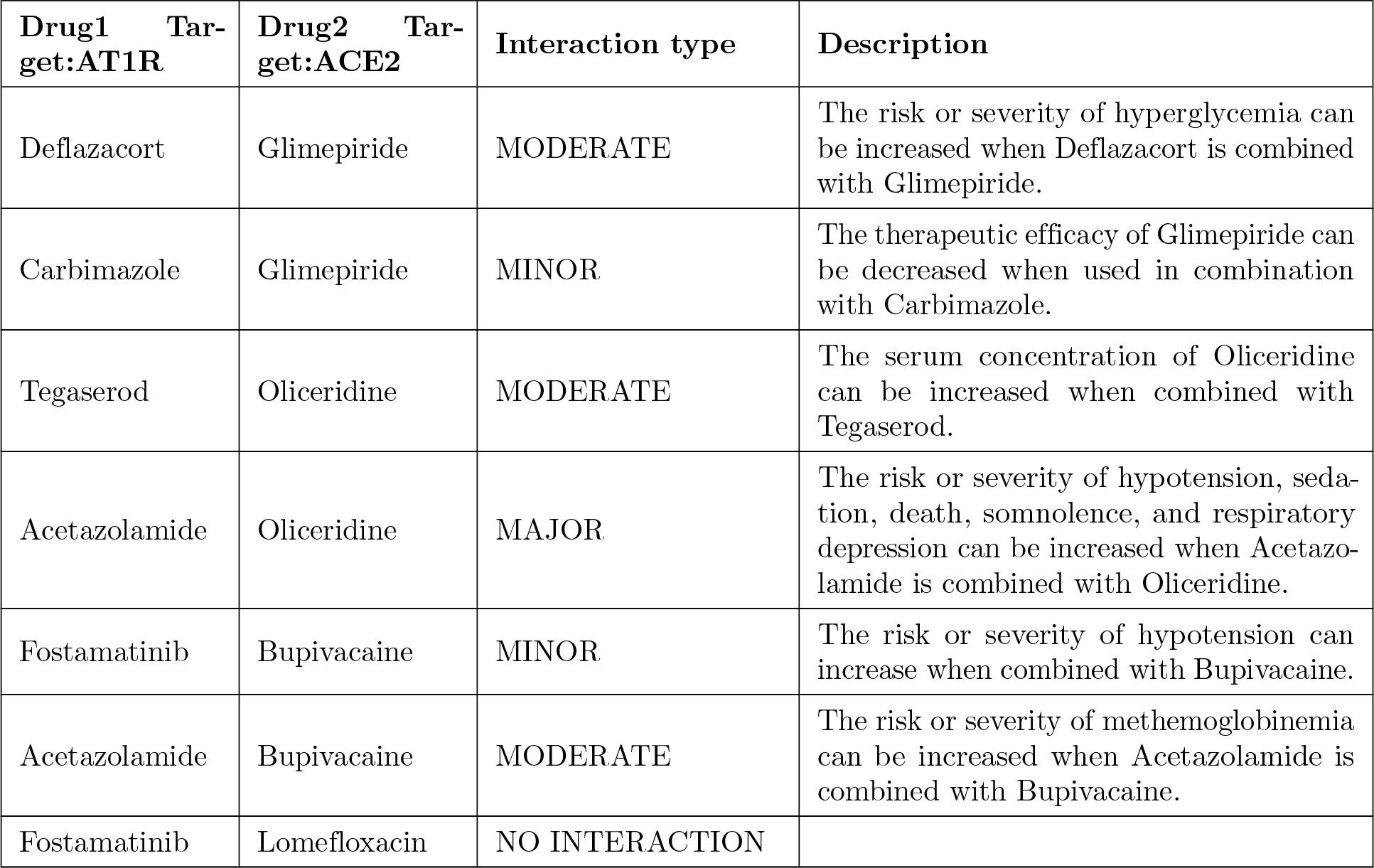
This table illustrates the drug-drug interaction between the drugs identified from the aforementioned DDI technique. The list of drugs in the first column is associated with AT1R inhibition, whereas the drugs in the second column are associated with ACE2 activation. (Source: DrugBank ^13^).

## 3. Results and Discussion

In our study, we incorporated a range of pivotal regulatory molecules, including ACE2, AT1R, ERK, IL-18, IL-6, IL-1*β*, NF*κ*B, Nrf2, and IFN-I, within our signaling pathway framework. The aim was to scrutinize their roles in orchestrating the upsurge of proinflammatory cytokines during the cytokine storm observed in severe COVID-19 patients. Visual representations of these participating signaling molecules in normal, infected, and drug-targeted scenarios have been presented in Figures 3-5. Quantifying these signaling molecules’ normalized expression profiles over time constitutes our network’s outcome. In this measurement, an expression level 1 denotes the peak relative expression of these molecules in a COVID-19 patient. In contrast, a level of 0 reflects an average individual’s minimal or basal expression.

**Figure 3.**
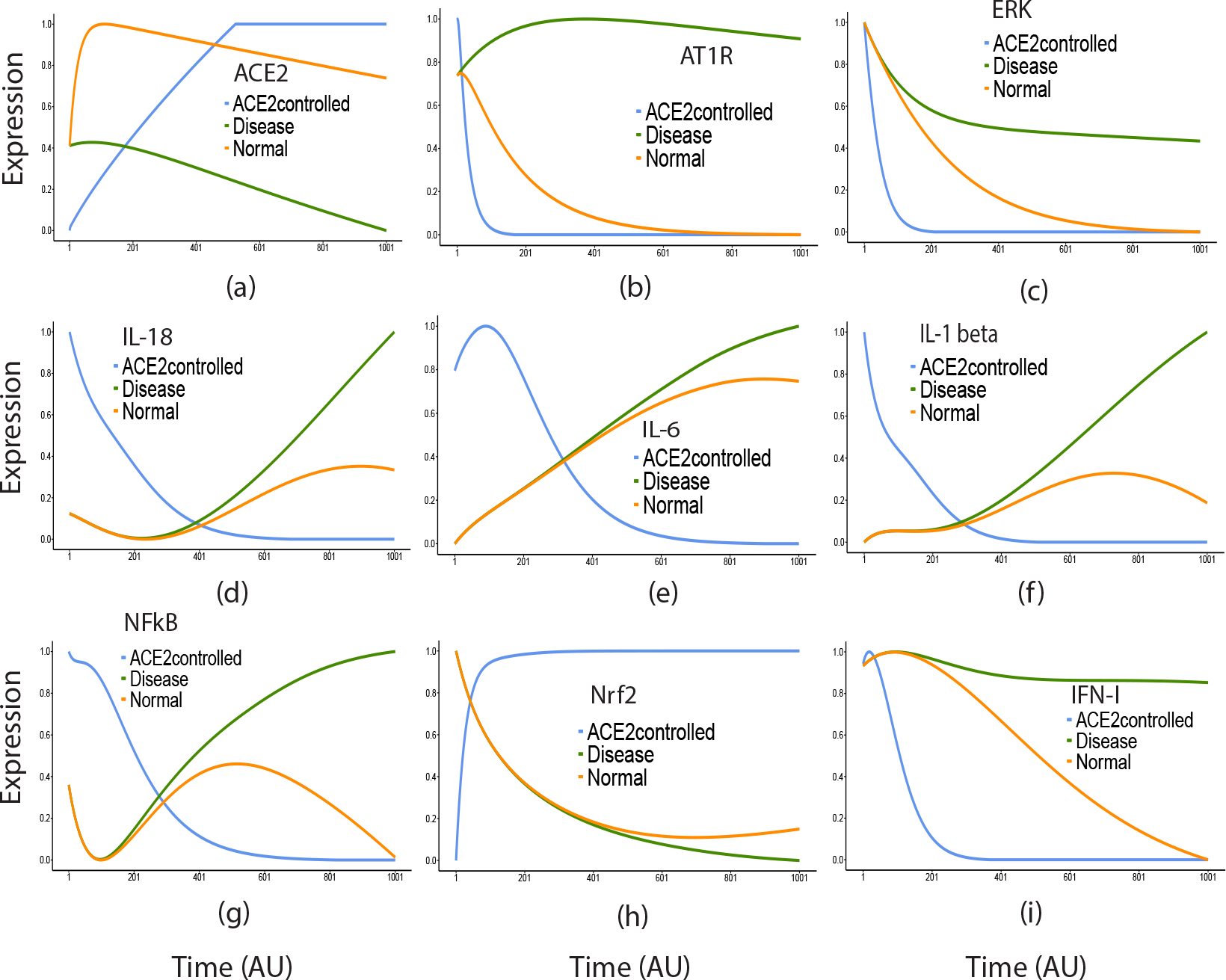
This figure illustrates a comparative analysis of the impact of ACE2 activation with normal and diseased conditions on nine key regulatory molecules. The analysis highlights ACE2 activation as a potential drug target.

Our results, depicted in Figure 3, revealed an increase in the expression of all signaling molecules within the disease system over time. An exception to this trend was observed with Nrf2, which exhibited consistently elevated expressions in both regular and disease systems. However, within the disease context, the Nrf2 levels experienced a slight decline after a specific interval ^27^. Remarkably, the elevation in the expression of these molecules closely paralleled the clinical observations concerning COVID-19 patients ^28^. For instance, Lebedeva et al. ^29^ conducted a comprehensive cytokine profiling of COVID-19 patients undergoing Tocilizumab therapy, uncovering statistically significant differences in normalized mRNA fluorescence values for IL-6, IL-1RA, IL-10, and G-CSF between patients with and without clinical endpoints.

Furthermore, our study underscored the significance of ACE2 expression, particularly in the context of SARS-CoV-2 entry. As the virus bound to the ACE2 receptor and underwent endocytosis, ACE2 expression assumed a pivotal role in evaluating the pathology of COVID-19. The endocytic process led to the downregulation of surface ACE2, resulting in unhindered Ang II accumulation. Given the presence of two ACE2 isoforms—one soluble (sACE2) and the other cell-specific—the broad expression of ACE2 encompassed tissues like the heart, kidneys, and lungs. Our data aligned with this pattern illustrated the release of ACE2 from the cytoplasmic membrane to the plasma in damaged tissues.

Consequently, the plasma ACE2 levels escalated in individuals afflicted with COVID-19. Conversely, cell type-specific expression in other tissues revealed that ACE and ACE2 protein expressions showcased higher ACE levels in cellular compartments, correlated with diminished ACE2 expression. This scenario accorded with our data, as depicted in Figure 3a.

Beyond the overall upregulation of proteins associated with signaling pathways, our observations regarding Nrf2 deviated from the norm. Nrf2 activity often experiences dysregulation in disease states, including inflammatory conditions. Notably, Nrf2 deficiency was linked to heightened ACE2 levels, while its activator contributed to the reduction of ACE2 levels ^30,31^. This phenomenon hints at the potential of NRF2 activation to diminish the availability of ACE2 for SARS-CoV-2 cell entry. Consequently, modulation of NRF2 emerges as a strategic avenue to mitigate the potential for SARS-CoV-2 infection (Figure 3).

Recent evidence has highlighted that SARS-CoV-2’s interaction with the ACE2 receptor prompts ACE2 downregulation, consequently stimulating AT1R ^32^. This evidence motivated us to investigate AT1R’s role as a potential drug target. As detailed earlier, we employed two distinct PID controllers to independently inhibit ACE2 and AT1R expression. Interestingly, our results showed that both interventions yielded comparable effects on the essential proteins associated with cytokine storms. Figure 4 illustrates the outcomes of AT1R inhibition. A comparative analysis of both targets in Figure 5 reveals significant effect similarity, albeit with slight distinctions in the expressions of NF*κ*B and IL-6 when AT1R was targeted. Based on these findings, we can infer that ACE2 holds promise as the primary drug target for addressing cytokine storms, whereas AT1R emerges as a viable alternative drug target to ACE2. Experts must predict potential drug-drug interactions in cases necessitating the simultaneous targeting of both ACE2 and AT1R. To address this, we conducted a rigorous analysis of the DrugBank database, yielding a comprehensive list of possible drug (for activation of ACE2)-drug (for inhibition of AT1R) interactions. As showcased in Table 3, these interactions have been stratified into three categories: “minor”, “moderate”, and “major”. However, the details of all probable interactions between ACE2 and AT1R targeting drugs have been presented in Supplementary Table S8.

**Figure 4.**
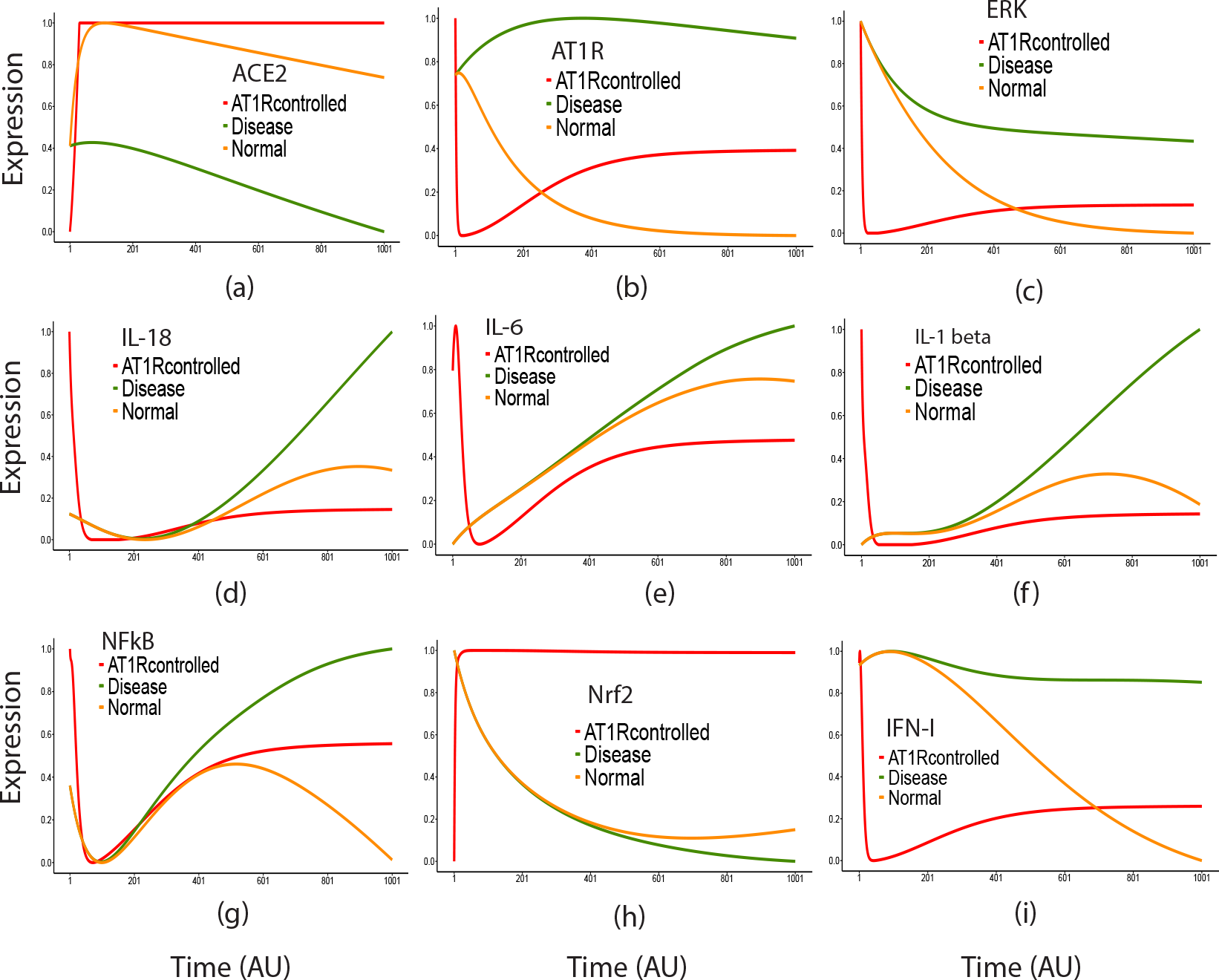
This figure illustrates a comparative analysis of the effects of AT1R inhibition on nine key regulatory molecules. The analysis indicates that AT1R inhibition can be an alternate strategy for considering as a potential drug target.

**Figure 5.**
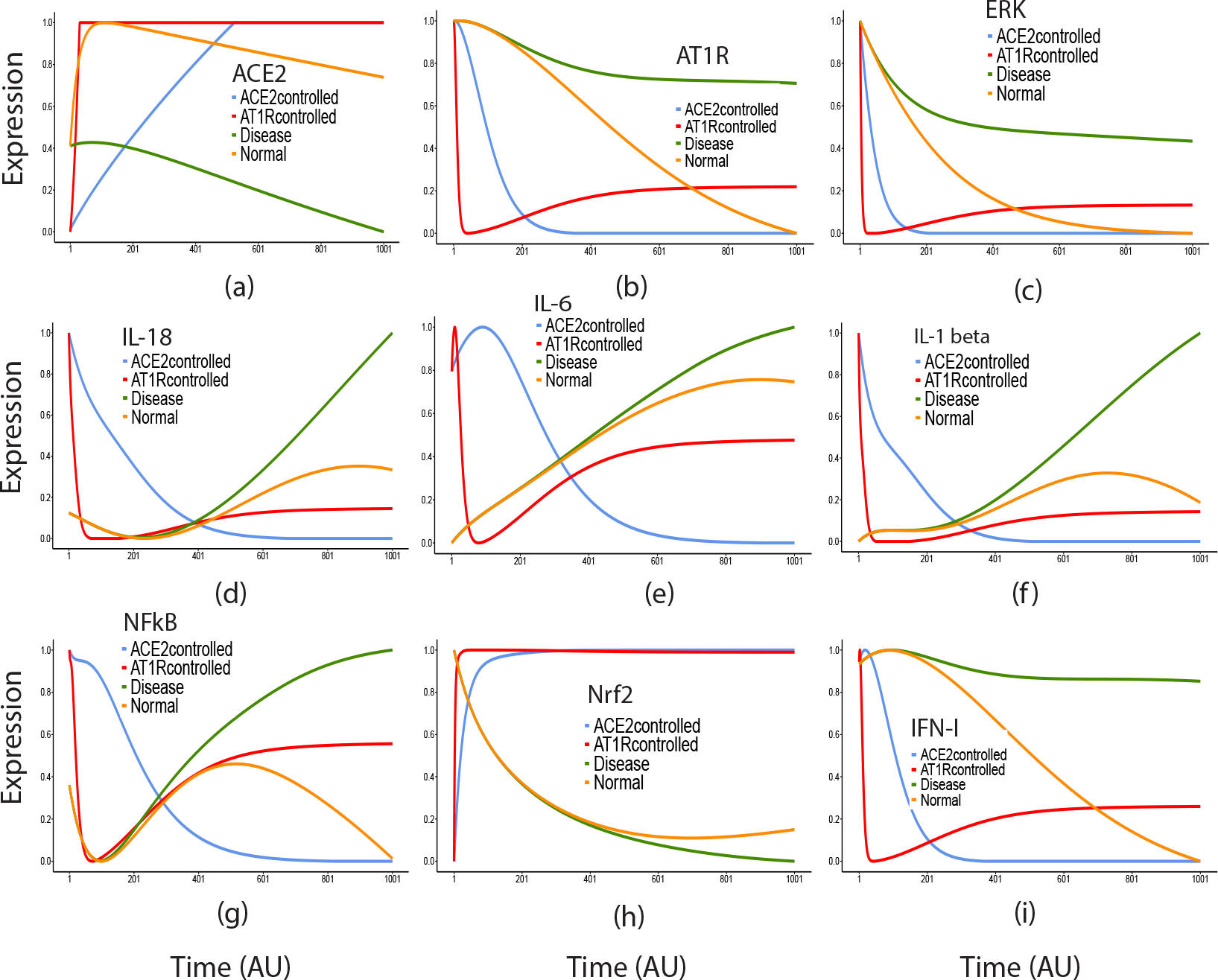
This figure presents a comparative analysis of the effects of ACE2 activation and AT1R inhibition on nine key regulatory molecules. The analysis demonstrates that ACE2 activation can be regarded as a potential drug target. The figure further illustrates that both ACE2 activation and AT1R inhibition have a similar effect on the key molecules of interest. Such observation implies that AT1R inhibition may serve as an alternative drug target. The effects of both interventions are compared under normal and diseased conditions.

Such tables also offer insights into the risks of a particular drug-drug interaction (Source: DrugBank^13^). Here, from Table 3, one can interpret that combining Deflazacort and Glimepiride presents a moderate risk of hyperglycemia. At the same time, Carbimazole’s interaction with Glimepiride slightly reduces the latter’s therapeutic effectiveness (minor). On the other hand, the combination of Acetazolamide and Oliceridine carries significant risks, including hypotension, sedation, respiratory depression, drowsiness, and, in severe cases, death (major). Fortunately, no significant interaction was observed between Fostamatinib and Lomefloxacin (no interaction). These insights are crucial for healthcare providers, emphasizing the need to balance therapeutic benefits and potential risks in medication management for patient safety and well-being.

Following training our deep learning model with a subset of ZINC15 drugs, we began a screening endeavor to repurpose these drugs during a crisis. This assessment involved testing diverse medicines, including antibiotics, antirheumatic drugs, peripheral nerve blockers, and angiotensin receptor blockers. Utilizing SWISSDOCK for docking experiments, we pinpointed the potential binding site in proximity to the C-terminus of the Receptor Binding Domain (RBD), as illustrated in Figure 6. Among the drugs considered in our study (Figure 6a), the protein-drug interactions displayed stabilities ranging from -5.96 to -9.47 Kcal/mol. Notably, Lomefloxacin, a fluoroquinolone antibiotic employed to treat bacterial infections such as bronchitis and urinary tract infections, emerged as the most promising candidate, demonstrating the highest binding affinity and stability ^33^. Its interaction was marked by electrostatic engagements with lysine74, asparagine102, and asparagine117 of the RBD, as depicted in Figure 6b.

**Figure 6.**
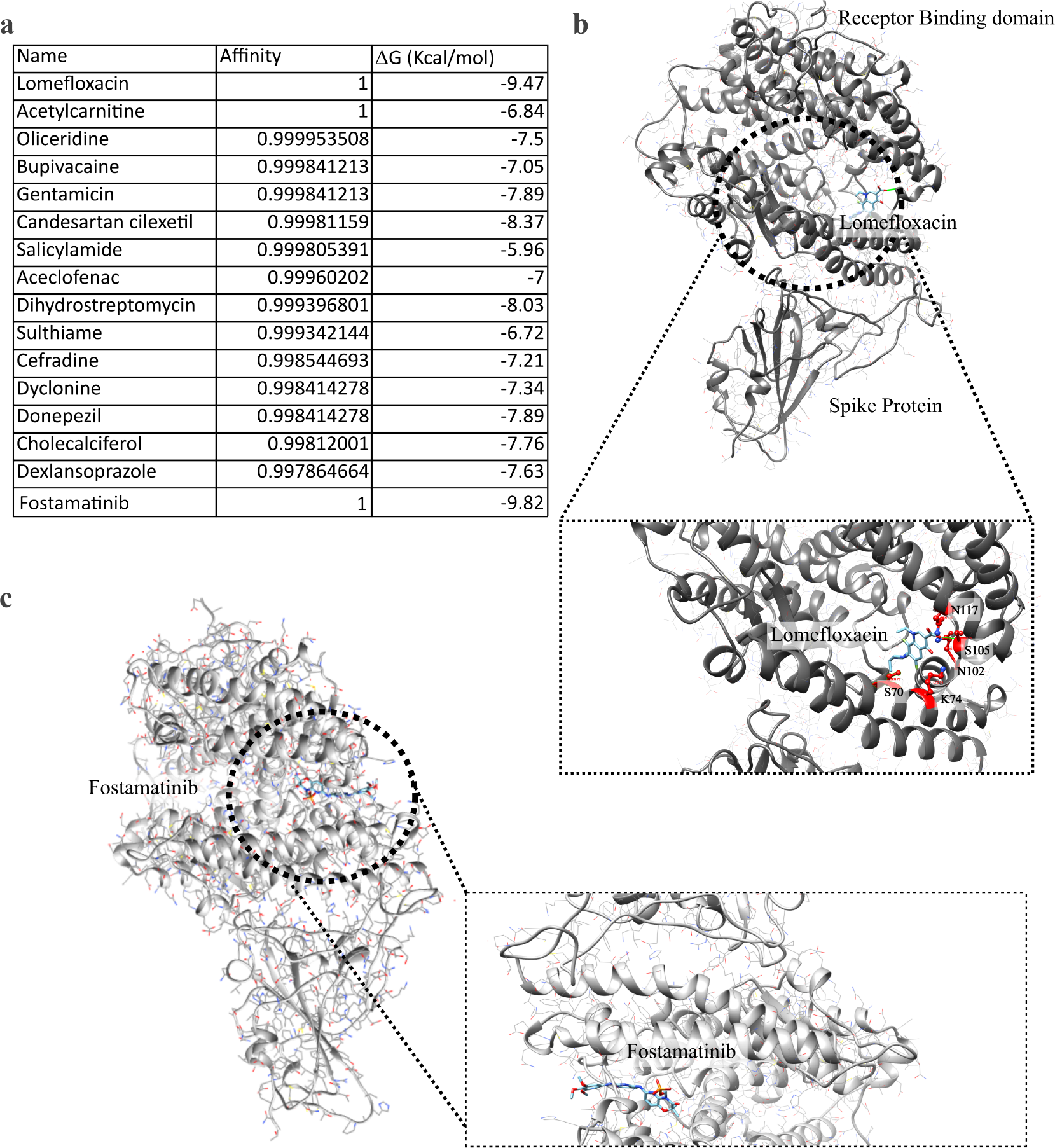
Molecular docking of drugs on RBD domain. a) Table summarizing the mathematical affinity and free energy of binding between the drug and RBD domain. The ΔG was calculated in Kcal/mol. b) Docking of Lomefloxacin with RBD domain of SARS-COV2 (PDB ID: 7DQA). Inset suggested the close interaction with the residues in the RBD domain. c) Docking of Fostamatinib with RBD domain of SARS-COV2 (PDB ID: 7DQA). Inset suggested the close interaction with the residues in the RBD domain.

Another hit, spleen tyrosine kinase inhibitor, Fostamatinib shows an edge over the other drugs to target ACE 2 protein in terms of thermodynamic stability. The stability of the Fostamatinib-ACE2 complex obtained from SWISSDOCK is -9.82 Kcal/mol. The aromatic moiety of the drug makes pi-pi interaction with the tryptophan (W) residue (position of the residue) due to its proximity. At the same time, it forms a cation-pi interaction with positively charged residues like lysine (K) and arginine (R) around the drug (Figure 6c). The stability of these two drugs can be justified by the interaction, as mentioned earlier, experienced by these drugs on the ACE 2 protein. It was observed that most of the *π*-*π* interactions have an energy in the range -0.5 to -2.0 kcal *mol*^*−*1^, while the cation–*π* interactions contribute an energy in the range -2 to -4 kcal *mol*^*−*1^. Further, with the increment of cations–*π* interaction pairs, the energy ranges from -6 to -13 kcal *mol*^*−*1^ ^34^. Moreover, our examination of potential interactions between Lomefloxacin and Fostamatinib did not uncover any noteworthy concerns. This finding reinforces the safety of co-administering these medications during cytokine storms (CS).

## 4. Conclusion

The relentless surge in human-to-human transmission of COVID-19 has proven exceedingly challenging to control, especially in developed countries with strained healthcare systems. Adding to the complexity is the resurgence of the epidemic, with instances of recovered patients testing positive again. The scientific community posits two main arguments to explain this phenomenon. First, the notion of “short memory” immunity suggests that immunity against COVID-19 could be temporary, fading after days or months. This could potentially render the virus “biphasic”, lying dormant before resurfacing with new symptoms. The second argument pertains to disease caused by distinct strains of the same virus. Such a study implies that an individual immune to one strain might remain vulnerable to a different strain of the same virus.

Amidst this backdrop, contemporary *in silico* investigations grounded in artificial intelligence (AI) have introduced a comprehensive pipeline for identifying a list of potential drugs to mitigate cytokine storms (CS), a pivotal concern in curbing the mortality rate associated with COVID-19 infections. In essence, managing the inflammatory response emerges as a critical aspect parallel to virus targeting. Approaches that simultaneously inhibit viral infection and regulate malfunctioning immune responses could synergistically combat pathologies at multiple stages. Simultaneously, the intricate interplay between immune dysfunction and the severity of COVID-19 underscores the need for prudence in vaccine development and evaluation.

Furthermore, there’s a pressing need for more detailed investigations utilizing AI-based models that consider specific protein-protein interactions and cross-talk within the host immune response to SARS-CoV-2. Such inquiries could unravel the determinants underlying healthy versus aberrant outcomes. This endeavor also holds promise in identifying biomarkers delineating immune correlates of protection and disease severity, facilitating effective patient triage.

Adding another layer of complexity, searching for an efficient drug remains challenging. The efficiency of a drug hinges on a delicate balance between strong affinity and stability. While strong affinity coupled with low stability ensures prompt dissociation after fulfilling its purpose, slow-release drugs necessitate both strong affinity and higher stability. Affinity is propelled by features like charge density and hydropathy, the number of interaction sites, and the alignment of the drug’s structure within the binding pocket. On the other hand, stability hinges on the energetic interactions and steric considerations imposed by the neighboring side chains around the binding sites. Consequently, long-term drug discovery efforts are dedicated to refining a more efficient deep learning-based model that optimizes these parameters, ultimately guiding the selection of the most suitable candidates based on specific requirements. Further, there is a lot of scope for optimization in the tuning of protein-drug interaction as the only strong binding would not be a determining factor for the potential drugs, fine-tuning is required

## Acknowledgement

We acknowledge the project with principal investigator Rajat K. De, sanctioned under MATRICS Special call under COVID-19 (File Number: MSC/2020/000350) in the Indian Statistical Institute by the Science and Engineering Research Board (SERB) under the Department of Science and Technology (DST), Government of India.

## Authors’ Contributions

AD and AB conceptualized the basic idea of PID controller-based pathway modeling for therapeutic intervention on cytokine storm in COVID-19. AB formulated the computational methodology and implemented it. KG performed docking. SB analyzed and validated the results. AD, AC, and RKD gave crucial theoretical input. AB, AD, SB, and KG wrote the first draft of the manuscript. AD, SB, RKD, and AC corrected it. AD and RKD supervised the entire work.

