## Supplementary Text for "In Silico Therapeutic Intervention on Cytokine Storm in COVID-19"

### Supplementary Material

##### Abstract

The recent global COVID-19 outbreak, attributed by the World Health Organization to the rapid spread of the severe acute respiratory syndrome coronavirus-2 (SARS-CoV-2), underscores the need for an extensive exploration of virological intricacies, fundamental pathophysiology, and immune responses. This investigation is vital to unearth potential therapeutic avenues and preventive strategies. Our study delves into the intricate interaction between SARS-CoV-2 and the immune system, coupled with exploring therapeutic interventions to counteract dysfunctional immune responses like the 'cytokine storm' (CS), a driver of disease progression. Understanding these immunological dimensions informs the design of precise multiepitope-targeted peptide vaccines using advanced immunoinformatics and equips us with tools to confront the cytokine storm. Employing a control theory-based approach, we scrutinize the perturbed behavior of key proteins associated with cytokine storm during COVID-19 infection. Our findings support ACE2 activation as a potential drug target for CS control and confirm AT1R inhibition as an alternative strategy. Leveraging deep learning, we identify potential drugs to individually target ACE2 and AT1R, with Lomefloxacin and Fostamatinib emerging as standout options due to their close interaction with ACE2. Their stability within the protein-drug complex suggests superior efficacy among many drugs from our deep-learning analysis. Moreover, there is a significant scope for optimization in fine-tuning protein-drug interactions. Strong binding alone may not be the sole determining factor for potential drugs; precise adjustments are essential. The application of advanced computational power offers novel solutions, circumventing time-consuming lab work. In scenarios necessitating both ACE2 and AT1R targeting, optimal drug combinations can be derived from our analysis of drug-drug interactions, as detailed in the manuscript.

$ Equal contribution
* Correspondence should be addressed to Rajat K. De, Ph.D., Machine Intelligence Unit, Indian Statistical Institute, 203 B.T. Road, Kolkata, West Bengal 700108, India, or Abhijit Dasgupta, Ph.D., Systems Biology Ireland, University College Dublin, Dublin 4, Ireland,.

**Table S1. It contains the list of ODEs representing the dynamics of the signaling pathway of COVID-19 considered in the main manuscript.**

1. $\dot{ANG-I} = -k_1 * [ANG-I] * (1 + F_1 * [ACE]) * (1 + F_2 * ACE2)/COVID + lamda_{ANG-I}$
2. $\dot{ANG-II} = -k_2 * [ANG-II] * (1 + F_3 * [ACE2])/COVID + k_3 * ANG-I * (1 + F_4 * ACE)$
3. $\dot{ANG-I-VII} = -k_4 * [ANG-I-VII] + k_5 * [ANG-II] * (1 + F_5 * ACE2)/COVID + k_6 * ANG-I-IX * (1 + F_6 * ACE)$
4. $\dot{ANG-I-IX} = -k_7 * [ANG-I-IX] + k_8 * [ANG-I] * (1 + F_7 * ACE2)/COVID$
5. $\dot{AT1R} = -k_9 * [AT1R]/(1 + F_8 * [ACE2]) + k_{10} * [ANG-II]$
6. $\dot{MASR} = -k_{11} * [MASR] + k_{12} * [ANG-I-VII]$
7. $\dot{ACE} = -k_{13} * [ACE] + lamda_{ACE}$
8. $\dot{ACE2} = -k_{14} * [ACE2] * COVID + lamda_{ACE2} * [ANG-I]$
9 $\dot{PLC} = -k_{15} * [PLC] + k_{16} * [AT1R]$
10. $\dot{PKC} = -k_{17} * [PKC] + k_{18} * [PLC]$
11. $\dot{TRAF6} = -k_{19} * [TRAF6] + k_{20} * [PKC] + k_{21} * [IL18] * [IL1b]$
12. $\dot{TAK1} = -k_{22} * [TAK1] + k_{23} * TRAF6$
13. $\dot{IKKb} = -k_{24} * [IKKb] + k_{25} * [TAK1] + k_{26} * [RSK]$
14. $\dot{NFkB} = -k_{27} * [NFkB] * (1 + F_9 * [Nrf2]) + k_{28} * [IKKb] + k_{29} * [IKKa] + k_{30} * [TRIF]$
15. $\dot{TLR3} = -k_{31} * [TLR3] + k_{32} * [AT1R]$
16. $\dot{TRIF} = -k_{33} * [TRIF] + k_{34} * [TLR3]$
17. $\dot{TLR78} = -k_{35} * [TLR78] + k_{36} * [AT1R]$
18. $\dot{MyD88} = -k_{37} * [MyD88] + k_{38} * [TLR78]$
19. $\dot{IL6} = -k_{39} * [IL6] + k_{40} * [NFkB] + k_{41} * [AP1]$
20. $\dot{IFN_I} = -gamma * [IFN_I] + k_{42} * [MyD88]$
21. $\dot{ERK} = -k_{43} * [ERK] + k_{44} * [AT1R]$
22. $\dot{RSK} = -k_{45} * [RSK] + k_{46} * [ERK]$
23. $\dot{PI3K} = -k_{47} * [PI3K] + k_{48} * [AT1R] + k_{49} * [IL18] * [IL1b] + k_{50} * [MyD88]$
24. $\dot{AKT} = -k_{51} * [AKT] + k_{52} * PI3K$
25. $\dot{Nrf2} = -k_{53} * [Nrf2] + k_{54}/(1 + F_{10} * [NFkB] * [AKT])$
26. $\dot{Caspase8} = -gamma * [Caspase8] + k_{55} * [MyD88]$
27. $\dot{AP1} = -k_{56} * [AP1] + k_{57} * [TAK1]$
28. $\dot{IL18} = -k_{58} * [IL18] + k_{59} * [PRO-IL-18] * (1 + F_{11} * [COMPLEX]) * [NFkB]$
29. $\dot{IL1b} = -k_{60} * [IL1b] + k_{61} * [PRO-IL-1b] * (1 + F_{12} * [COMPLEX]) * [NFkB]$
30. $\dot{NLRP3} = -k_{62} * [NLRP3] + k_{63} * [NFkB]$
31. $\dot{PRO-IL-18} = -k_{64} * [PRO-IL-18] + k_{65} * [NFkB]$
32. $\dot{PRO-IL-1b} = -k_{66} * [PRO-IL-1b] + k_{67} * [NFkB]$
33. $\dot{PRO_CASPASE_1} = -k_{68} * [PRO_CASPASE_1] + k_{69} * [NFkB]$
34. $\dot{COMPLEX} = -k_{70} * [COMPLEX] + k_{71} * [NLRP3] * [PRO_CASPASE_1]/(1 + F_{13} * [Nrf2])$
35. $\dot{IKKa} = -k_{72} * [IKKa] + k_{73} * [AKT] * [TRAF6]$

**Table S2. IF-THEN rule base for FIS to capture the fuzzy relationship between AKT, NF$\kappa$B & Nrf2**

| **IF (AKT & NF$\kappa$B)** | | **Then (Nrf2)** |
|---|---|---|
| $AKT$ | $NF\kappa B$ | $Nrf2$ |
| **LOW** | **LOW** | **LOW** |
| **LOW** | **MEDIUM** | **LOW** |
| **LOW** | **HIGH** | **LOW** |
| **MEDIUM** | **LOW** | **MEDIUM** |
| **MEDIUM** | **MEDIUM** | **MEDIUM** |
| **MEDIUM** | **HIGH** | **LOW** |
| **HIGH** | **LOW** | **HIGH** |
| **HIGH** | **MEDIUM** | **MEDIUM** |
| **HIGH** | **HIGH** | **MEDIUM** |

**Table S3. IF-THEN rule base for FIS to capture the fuzzy relationship between ANG-II, ACE2 & AT1R, MASR**

| IF (ANG-II & ACE2) | | Then (AT1R & MASR) | |
|---|---|---|---|
| *ANG − II* | *ACE2* | *AT1R* | *MASR* |
| LOW | LOW | LOW | LOW |
| LOW | MEDIUM | LOW | MEDIUM |
| LOW | HIGH | LOW | MEDIUM |
| MEDIUM | LOW | MEDIUM | LOW |
| MEDIUM | MEDIUM | MEDIUM | MEDIUM |
| MEDIUM | HIGH | LOW | MEDIUM |
| HIGH | LOW | HIGH | LOW |
| HIGH | MEDIUM | HIGH | MEDIUM |
| HIGH | HIGH | LOW | HIGH |

**Table S4. IF-THEN rule base for FIS to capture fuzzy relationship between NF$\kappa$B, AP1 & IL6, IL18, IL1$\beta$.**

| IF (NF$\kappa$B & AP1) | | Then (IL6 & IL18 & IL1$\beta$) | | |
|---|---|---|---|---|
| $NF\kappa B$ | $AP1$ | $IL6$ | $IL18$ | $IL1\beta$ |
| LOW | LOW | LOW | LOW | LOW |
| LOW | MEDIUM | MEDIUM | LOW | LOW |
| LOW | HIGH | MEDIUM | LOW | LOW |
| MEDIUM | LOW | MEDIUM | MEDIUM | MEDIUM |
| MEDIUM | MEDIUM | MEDIUM | MEDIUM | MEDIUM |
| MEDIUM | HIGH | HIGH | MEDIUM | MEDIUM |
| HIGH | LOW | MEDIUM | MEDIUM | MEDIUM |
| HIGH | MEDIUM | HIGH | MEDIUM | MEDIUM |
| HIGH | HIGH | HIGH | HIGH | HIGH |

**Table S5. IF−THEN rule base for FIS to capture fuzzy relationship between AT1R & TLR3 & TLR7/8 and NF$\kappa$B**

| IF (AT1R & TLR3 & TLR7/8) | | | Then (NF$\kappa$B) |
|---|---|---|---|
| AT1R | TLR3 | TLR7/8 | NF$\kappa$B |
| LOW | LOW | LOW | LOW |
| LOW | LOW | MEDIUM | MEDIUM |
| LOW | LOW | HIGH | MEDIUM |
| LOW | MEDIUM | LOW | MEDIUM |
| LOW | MEDIUM | MEDIUM | MEDIUM |
| LOW | MEDIUM | HIGH | HIGH |
| LOW | HIGH | LOW | MEDIUM |
| LOW | HIGH | MEDIUM | HIGH |
| LOW | HIGH | HIGH | HIGH |
| MEDIUM | LOW | LOW | LOW |
| MEDIUM | LOW | MEDIUM | MEDIUM |
| MEDIUM | LOW | HIGH | HIGH |
| MEDIUM | MEDIUM | LOW | MEDIUM |
| MEDIUM | MEDIUM | MEDIUM | MEDIUM |
| MEDIUM | MEDIUM | HIGH | HIGH |
| MEDIUM | HIGH | LOW | HIGH |
| MEDIUM | HIGH | MEDIUM | HIGH |
| MEDIUM | HIGH | HIGH | HIGH |
| HIGH | LOW | LOW | HIGH |
| HIGH | LOW | MEDIUM | HIGH |
| HIGH | LOW | HIGH | HIGH |
| HIGH | MEDIUM | LOW | HIGH |
| HIGH | MEDIUM | MEDIUM | HIGH |
| HIGH | MEDIUM | HIGH | HIGH |
| HIGH | HIGH | LOW | HIGH |
| HIGH | HIGH | MEDIUM | HIGH |
| HIGH | HIGH | HIGH | HIGH |

**Table S6. IF−THEN rule base for FIS to capture fuzzy relationship between TRAF6 & PI3K & TAK1 and IKK$\alpha$**

| **IF (TRAF6 & PI3K & TAK1)** | | | **Then (IKK$\alpha$)** |
|---|---|---|---|
| **TRAF6** | **PI3K** | **TAK1** | **IKK$\alpha$** |
| **LOW** | **LOW** | **LOW** | **LOW** |
| **LOW** | **LOW** | **MEDIUM** | **LOW** |
| **LOW** | **LOW** | **HIGH** | **LOW** |
| **LOW** | **MEDIUM** | **LOW** | **MEDIUM** |
| **LOW** | **MEDIUM** | **MEDIUM** | **MEDIUM** |
| **LOW** | **MEDIUM** | **HIGH** | **HIGH** |
| **LOW** | **HIGH** | **LOW** | **MEDIUM** |
| **LOW** | **HIGH** | **MEDIUM** | **MEDIUM** |
| **LOW** | **HIGH** | **HIGH** | **HIGH** |
| **MEDIUM** | **LOW** | **LOW** | **LOW** |
| **MEDIUM** | **LOW** | **MEDIUM** | **LOW** |
| **MEDIUM** | **LOW** | **HIGH** | **LOW** |
| **MEDIUM** | **MEDIUM** | **LOW** | **MEDIUM** |
| **MEDIUM** | **MEDIUM** | **MEDIUM** | **MEDIUM** |
| **MEDIUM** | **MEDIUM** | **HIGH** | **MEDIUM** |
| **MEDIUM** | **HIGH** | **LOW** | **HIGH** |
| **MEDIUM** | **HIGH** | **MEDIUM** | **HIGH** |
| **MEDIUM** | **HIGH** | **HIGH** | **HIGH** |
| **HIGH** | **LOW** | **LOW** | **MEDIUM** |
| **HIGH** | **LOW** | **MEDIUM** | **MEDIUM** |
| **HIGH** | **LOW** | **HIGH** | **MEDIUM** |
| **HIGH** | **MEDIUM** | **LOW** | **HIGH** |
| **HIGH** | **MEDIUM** | **MEDIUM** | **HIGH** |
| **HIGH** | **MEDIUM** | **HIGH** | **HIGH** |
| **HIGH** | **HIGH** | **LOW** | **HIGH** |
| **HIGH** | **HIGH** | **MEDIUM** | **HIGH** |
| **HIGH** | **HIGH** | **HIGH** | **HIGH** |

**Table S7.** This table illustrates the estimated parameter values used in the present investigation.

| Constant type | Parameter name | Estimated value |
|---|---|---|
| Binding rate constant | $k_1$ | 0.174636052 |
| | $k_2$ | 0.039401179 |
| | $k_3$ | 0.010945041 |
| | $k_4$ | 0.955654623 |
| | $k_5$ | 0.203502429 |
| | $k_6$ | 0.438953835 |
| | $k_7$ | 0.570824453 |
| | $k_8$ | 0.380566733 |
| | $k_9$ | 0.999856658 |
| | $k_{10}$ | 0.947392098 |
| | $k_{11}$ | 0.938383778 |
| | $k_{12}$ | 0.690134295 |
| | $k_{13}$ | 0.385986282 |
| | $k_{14}$ | 0.010318883 |
| | $k_{15}$ | 0.953398619 |
| | $k_{16}$ | 0.764546773 |
| | $k_{17}$ | 0.74971923 |
| | $k_{18}$ | 0.134124119 |
| | $k_{19}$ | 0.211346054 |
| | $k_{20}$ | 0.272183699 |
| | $k_{21}$ | 0.627232545 |
| | $k_{22}$ | 0.818971807 |
| | $k_{23}$ | 0.626451543 |
| | $k_{24}$ | 0.81083752 |
| | $k_{25}$ | 0.533736501 |
| | $k_{26}$ | 0.674500588 |
| | $k_{27}$ | 0.941075388 |
| | $k_{28}$ | 0.453539901 |
| | $k_{29}$ | 0.132941664 |
| | $k_{30}$ | 0.383559342 |
| | $k_{31}$ | 0.191121776 |
| | $k_{32}$ | 0.900747028 |
| | $k_{33}$ | 0.218097136 |
| | $k_{34}$ | 0.193334346 |
| | $k_{35}$ | 0.580831194 |
| | $k_{36}$ | 0.475165235 |
| | $k_{37}$ | 0.806736351 |
| | $k_{38}$ | 0.528762003 |
| | $k_{39}$ | 0.432898 |
| | $k_{40}$ | 0.443234847 |
| | $k_{41}$ | 0.126080352 |
| | $k_{42}$ | 0.658288913 |
| | $k_{43}$ | 0.638269164 |
| | $k_{44}$ | 0.220474924 |
| | $k_{45}$ | 0.203223944 |
| | $k_{46}$ | 0.136014375 |
| | $k_{47}$ | 0.187727885 |
| | $k_{48}$ | 0.219760727 |
| | $k_{49}$ | 0.950841183 |
| | $k_{50}$ | 0.214200254 |
| | $k_{51}$ | 0.614056522 |
| | $k_{52}$ | 0.306453913 |
| | $k_{53}$ | 0.751397309 |
| | $k_{54}$ | 0.416853981 |
| | $k_{55}$ | 0.525199213 |
| | $k_{56}$ | 0.382271904 |
| | $k_{57}$ | 0.573078359 |
| | $k_{58}$ | 0.223545642 |
| | $k_{59}$ | 0.282253756 |
| | $k_{60}$ | 0.547590367 |
| | $k_{61}$ | 0.875305673 |
| | $k_{62}$ | 0.97787478 |
| | $k_{63}$ | 0.968650531 |
| | $k_{64}$ | 0.340662922 |
| | $k_{65}$ | 0.492791958 |
| | $k_{66}$ | 0.454816343 |
| | $k_{67}$ | 0.510962386 |
| | $k_{68}$ | 0.693713081 |
| | $k_{69}$ | 0.521764809 |
| | $k_{70}$ | 0.382326072 |
| | $k_{71}$ | 0.587123881 |
| | $k_{72}$ | 0.155295828 |
| | $k_{73}$ | 0.318293477 |
| Feedback constant | $F_1$ | 0.449045917 |
| | $F_2$ | 0.457456135 |
| | $F_3$ | 0.01761964 |
| | $F_4$ | 0.320045019 |
| | $F_5$ | 0.384961666 |
| | $F_6$ | 0.245422845 |
| | $F_7$ | 0.510526962 |
| | $F_8$ | 0.985280949 |
| | $F_9$ | 0.521553366 |
| | $F_{10}$ | 0.43802267 |

*Continued on next page*

Table S7 – *Continued from previous page*

| Constant type | Parameter name | Estimated value |
|---|---|---|
| | $F_{11}$ | 0.50758626 |
| | $F_{12}$ | 0.504041161 |
| | $F_{13}$ | 0.502185969 |
| Decay constant | $lamda_{ANG-I}$ | 0.918499208 |
| | $lamda_{ACE}$ | 0.54699201 |
| | $lamda_{ACE2}$ | 0.99980658 |
| | $gamma$ | 0.499749004 |

**Table S8.** This table illustrates the drug-drug interaction between the drugs identified from the DDI technique mentioned in the main manuscript. The list of drugs in the first column is associated with AT1R inhibition, whereas the drugs in the second column are associated with ACE2 activation. (Source: DrugBank[1]).

| Drug 1 | Drug 2 | Interaction type | Description |
|---|---|---|---|
| Ferrous fumarate | Lomefloxacin | MODERATE | Ferrous fumarate can cause a decrease in the absorption of Lomefloxacin, resulting in a reduced serum concentration and potentially a decrease in efficacy. |
| Ivabradine | Lomefloxacin | MODERATE | Ivabradine may increase the QTc-prolonging activities of Lomefloxacin. |
| Pramlintide | Lomefloxacin | MODERATE | The therapeutic efficacy of Pramlintide can be increased when combined with Lomefloxacin. |
| Deflazacort | Lomefloxacin | MODERATE | The risk or severity of tendinopathy can be increased when Deflazacort is combined with Lomefloxacin. |
| Tegaserod | Lomefloxacin | MODERATE | The metabolism of Lomefloxacin can be decreased when combined with Tegaserod. |
| Acetazolamide | Glimepiride | MODERATE | The therapeutic efficacy of Glimepiride can be increased when used in combination with Acetazolamide. |
| Glimepiride | Pramlintide | MODERATE | The risk or severity of hypoglycemia can be increased when Glimepiride is combined with Pramlintide. |
| Deflazacort | Glimepiride | MODERATE | The risk or severity of hyperglycemia can be increased when Deflazacort is combined with Glimepiride. |
| Carbimazole | Glimepiride | MINOR | The therapeutic efficacy of Glimepiride can be decreased when used in combination with Carbimazole. |
| Tegaserod | Oliceridine | MODERATE | The serum concentration of Oliceridine can be increased when combined with Tegaserod. |
| Acetazolamide | Oliceridine | MAJOR | The risk or severity of hypotension, sedation, death, somnolence, and respiratory depression can be increased when Acetazolamide is combined with Oliceridine. |
| Fostamatinib | Bupivacaine | MINOR | The risk or severity of hypotension can increase when combined with Bupivacaine. |
| Acetazolamide | Bupivacaine | MODERATE | The risk or severity of methemoglobinemia can be increased when Acetazolamide is combined with Bupivacaine. |
| Ripretinib | Bupivacaine | MODERATE | The risk or severity of methemoglobinemia can be increased when Ripretinib is combined with Bupivacaine. |
| Tegaserod | Bupivacaine | MODERATE | The metabolism of Bupivacaine can be decreased when combined with Tegaserod. |
| Fostamatinib | Candesartan cilexetil | MINOR | Fostamatinib may increase the antihypertensive activities of Candesartan cilexetil. |
| Acetazolamide | Caffeine / Salicylamide | MODERATE | Acetazolamide may increase the excretion rate of Caffeine which could result in a lower serum level and potentially a reduction in efficacy. |
| Ripretinib | Caffeine / Salicylamide | MODERATE | Caffeine may decrease the excretion rate of Ripretinib, which could result in a higher serum level. |
| Tegaserod | Caffeine / Salicylamide | MODERATE | Caffeine may decrease the excretion rate of Tegaserod, which could result in a higher serum level. |
| Deflazacort | Caffeine / Salicylamide | MINOR | The risk or severity of gastrointestinal irritation can be increased when Deflazacort is combined with Salicylamide. |
| Carbimazole | Caffeine / Salicylamide | MODERATE | Carbimazole may decrease the excretion rate of Caffeine, which could result in a higher serum level. |
| Acetazolamide | Aceclofenac | MODERATE | Acetazolamide may increase the excretion rate of Aceclofenac, which could result in a lower serum level and potentially a reduction in efficacy. |
| Deflazacort | Aceclofenac | MINOR | The risk or severity of gastrointestinal irritation can be increased when Deflazacort is combined with Aceclofenac. |
| Lomefloxacin | Fostamatinib | NO INTERACTION | |
